# Common core respiratory bacteriome of the blue whale *Balaenoptera musculus*, in the Gulf of California

**DOI:** 10.1101/2022.12.29.522252

**Authors:** Carlos A. Domínguez-Sánchez, Roberto C. Álvarez-Martínez, Diane Gendron, Karina Acevedo-Whitehouse

## Abstract

The number of strandings and unusual mortality events that involve cetaceans may have increased, and potential pathogens of the respiratory tract have been found during the examination of individuals in many of these events. However, investigating the health of free-ranging large whales is logistically complex. Given that the core microbiome is key to understanding host-bacteria relationships and to identifying their relevance for individual and population health, we characterized the core respiratory bacteriome of the Eastern North Pacific blue whale, *Balaenoptera musculus*, using blow samples collected by a small quadracopter drone. 16S rRNA gene high-throughput sequencing revealed 1,326 amplicon sequence variants (ASVs), of which 11 were shared by more than 50% of all blue whales and had a relative abundance higher than 0.02%. *Cutibacterium, Oceanivirga, Tenacibaculum*, and *Psychrobacter* composed the common core respiratory bacteriome of the blue whale. Additionally, compositional analysis identified 15 bacterial classes dominated by Gammaproteobacteria (27.14%), Bacteroidea (19.83%), and Clostridia (12.89%) as the most representative classes in the respiratory tract of blue whales. However, two whales had a high abundance of bacteria with pathogenic potential, namely M*ycoplasma* spp. and *Streptococcus* spp. in their blow. Both bacterial genera have been associated with pulmonary diseases in mammals. Ours is the first study to characterize the respiratory bacteriome of apparently healthy blue whales and is a baseline for future long-term studies on blue whale health, an endangered species of conservation concern.

## Introduction

The advent of modern technologies that allow the identification of all bacteria present in environmental or clinical samples (Haegeman et al., 2013; Salter et al., 2014; Rhodes et al., 2022) has led to a myriad of studies on the abundance, diversity, and structure of the microbiome of different species (Nelson et al., 2015; Watkins et al., 2022). One of the reasons why it is paramount to increase our knowledge about the microbiome of a given species is because the microbial communities associated with a particular epithelium can impact the host’s physiology (Foster et al., 2017), and even influence its health status (Huang, Yang, & Li, 2016; Bierlich et al., 2018; Panthee et al., 2022; Watkins et al., 2022). For example, respiratory infections can occur when opportunistic microorganisms, which are normally part of the microbiome of a healthy respiratory mucosa, preferentially flourish under certain conditions (Dickson et al., 2016; Huffnagle, Dickson, & Lukacs, 2017; Rhodes et al., 2022), and *de novo* infections may occur if individuals are exposed to pathogens. In turn, infections can trigger changes in the diversity and composition of the original microbial communities, an event known as dysbiosis (Gagliardi et al., 2018; Robles-Malagamba et al., 2020; Sehnal et al., 2021). As a result, the composition of the microbiome may be a better predictor of disease progression than the presence of a specific pathogen commonly associated with the disease. Thus, understanding the composition of the microbiome and how it differs between healthy and sick animals could become an important tool for assessing an individual’s health status (Shreiner, Kao, & Young, 2015). Such knowledge, in turn, is paramount for animal conservation efforts, as the impact of diseases on wild animal populations is increasingly becoming evident (Hu, Han, & He, 2023). This has led to a recent surge in studies that have begun to examine the microbiome of species of conservation concern (e.g. Dallas & Warne, 2023; Jiang et al., 2022; Vecchioni et al., 2022).

When attempting to use the microbiome to assess health status, commensal, opportunistic, and transient bacteria must be identified (Huang, Yang, & Li, 2016; Infante-Villamil, Huerlimann, & Jerry, 2021). To do this, it is necessary to identify the bacterial taxa that predominate in the community and are shared by healthy individuals (Huse et al., 2012; Willis, Bunge, & Whitman, 2017; Risely, 2020). This is known as the core microbiome and is well known to play an important role in maintaining functional stability and homeostasis of a specific habitat within the host (Hernandez-Agreda, Gates, & Ainsworth, 2017; Thomas et al., 2017; Björk et al., 2018; Ross, Rodrigues-Hoffmann, & Neufeld, 2019). The definition of the core microbiome varies across authors, although they tend to overlap in many of the components of the microbial community (Risely, 2020). Different approaches to define the core microbiome have included temporal stability (Ozkan et al., 2017), functional level (Dinsdale et al., 2008), ecological influence (Coyte et al., 2019), host fitness (Shapira, 2016), and bacterial occupancy frequency, which refers to the most common bacterial taxa that are shared by a considerable proportion of hosts (Ingala et al., 2018; Nishida & Ochman, 2018; Risely, 2020; Toro et al., 2021).

Identifying the common core bacteriome (i.e. the bacterial community of the microbiome) requires setting the detection threshold (relative abundance) and the percentage of occurrence (prevalence) of bacterial taxa (Astudillo-García et al., 2017), criteria which have varied widely among published studies (Risely, 2020), with common core annotations ranging from as low as 30% (e.g., Ainsworth et al., 2015) to 100% prevalence (e.g. (Huse et al., 2012; Apprill et al., 2017; Hernandez-Agreda, Gates, & Ainsworth, 2017; Antwis et al., 2018), and detection thresholds varying from 0.001% to 0.1% (Astudillo-García et al., 2017; Antwis et al., 2018). Given that biological justifications for such prevalence and threshold values are rare (Risely, 2020), it is important to recognize the arbitrary aspect of the common core definition and to be cautious when interpreting the results. However, the bacteriome common core tends to be robust despite varying definitions, particularly when samples from closely related individuals are analysed (Risely, 2020; Vendl et al., 2020).

Bacteria of the mammalian microbiome are found in composite communities (Lee & Mazmanian, 2010; Rhodes et al., 2022), whose diversity and abundance are determined by multiple interactions between species (Shade & Handelsman, 2012; Stubbendieck, Vargas-Bautista, & Straight, 2016). The cetacean microbiome has recently begun to be studied and important initial assessments of microbial diversity have been made for a few species (Johnson et al., 2009a; Hooper et al., 2019; Apprill et al., 2020; Van Cise et al., 2020; Toro et al., 2021; Rhodes et al., 2022). As long-lived cetaceans that contribute to the movement and storage of carbon in the ocean, baleen whales play a vital role in the marine ecosystem (Pershing et al., 2010), and are considered sentinels of ocean health (Sharp et al., 2019; Palmer et al., 2022). However, to date, little is known about the respiratory microbiome of baleen whales, and while some opportunistic pathogens of the respiratory tract have been described for live free-ranging individuals of a few baleen whale species (Acevedo-Whitehouse et al., 2010), to the best of our knowledge, there is only one published study on the core respiratory microbiome of a baleen whale, the humpback whale *Megaptera novaeangliae* (Apprill et al., 2017).

Blue whales are among the world’s largest and most iconic animals and play an essential role in marine ecosystems (Attard, Beheregaray, & Möller, 2016). Owing to their long migration and diverse habitats, they are also important indicators of the health of oceans worldwide (Rice et al., 2022). Studies on the respiratory microbiome of blue whales are crucial for understanding their health and well-being in their natural environment because the bacteria in their respiratory system can provide valuable information about their exposure to pathogens (Raverty et al., 2017). This can aid in identifying potential diseases and developing effective conservation strategies for marine mammals (Raverty et al., 2020), especially considering global and local environmental changes affecting oceans (Ferguson et al., 2021; Aguilar & Borrell, 2022; Barlow et al., 2023). Therefore, studying the respiratory microbiome is key to conserving this iconic species and protecting its long-term survival. In this study, we characterized the common core respiratory bacteriome of the Eastern North Pacific blue whale using next-generation sequencing on 17 blow samples collected from adult blue whales by a non-invasive drone-based technique (Domínguez-Sánchez et al., 2018) during the boreal winter months in the Gulf of California.

## Materials and Methods

### Sample collection

Using a small Phantom 3 quadracopter drone (DJI Innovations, China) with floaters and sterile Petri dishes, we collected 17 blow samples from 17 individual blue whales sampled between February and March 2016 and 2017 in Loreto Bay National Park (25°51’51’’N, 111°07’18’’O) within the Gulf of California, Mexico. The number of sampled whales represents 17% of the estimated 100 blue whales that reside during winter/spring in the southwestern Gulf of California (mark-recapture data from 1994-2006; Gendron & Ugalde de la Cruz, 2012; SEMARNAT, 2018). Each whale was photo-identified prior to sample collection (Gendron & Ugalde de la Cruz, 2012). The approach to the whale with the drone was made from the caudal fin heading towards the head to minimize disturbance, and sampling was conducted at a height between 3 and 4 m above the blowhole (Domínguez-Sánchez et al., 2018).

For each sample, the blow droplets were swabbed directly from the Petri dish using sterile cotton-tipped swabs. These were then transferred to a sterile 1.5 mL cryogenic microtube containing 500 µL of 96% molecular grade ethanol and kept frozen in a liquid nitrogen container until processing. To address potential contamination, all necessary precautions were taken, always including the use of sterile gloves and face masks during sample handling and processing. In addition, we included three technical controls:1) LabControl (DNA extraction from kit reagents), 2) HumanSneeze (human sneeze sampled from the person who collected and processed the samples), and 3) SeaWater (seawater collected at a depth of 0.10 m in the exact location where we sampled the whale blows).

### DNA extraction, PCR amplification and sequencing

Total DNA was isolated from the whale blow and technical control samples in one batch using a QIAamp DNA Mini Kit (QIAGEN, Germany). The primers used for sequencing the 16SrRNA V3 and V4 regions were 341F (5’-CCTACGGGNGGCWGCAG) and 785R (5’-GACTACHVGGGTATCTAATCC), which amplified a single product of approximately 460 bp (Thijs et al., 2017). The PCR program used an initial denaturation step at 95°C for 3 min; 25 cycles of 95°C for 30 s, 55°C for 30 s, and 72°C for 30s; and a final extension step at 72°C for 5 min. Each 25 µL-reaction contained 12.5 ng of extracted DNA, 5 µM of barcoded primers and 2x KAPA HiFi HotStart Ready Mix (KAPABIOSYSTEM, Cape Town, South Africa). 1 µl of each sample was run on a 2100 Bioanalyzer (Agilent Technologies, CA, USA) with an Agilent DNA 1000 chip (Agilent Technologies, CA, USA) to verify amplicon size. AMPure XP beads (New England BioLabs, USA) were used to remove unused primers and primer dimers. Amplicons were sequenced over 2- by 250-bp MiSeq at the Unit of Sequencing and Identification of Polymorphisms of the National Institute of Genomic Medicine (Instituto Nacional de Medicina Genómica, Unidad de Secuenciación e Identificación de Polimorfismos) in Mexico.

### 16S rRNA sequence data processing

Quality control overview was performed using FASTQC (Andrews, 2010). This allowed us to obtain a quick impression of the data and avoid downstream problems. The raw sequences were then imported into R v.4.2.1 (R Core Team, 2022), where all subsequent analyses were carried out. We used the Divisive Amplicon Denoising Algorithm 2 (*dada2*) v.1.26.0 (Callahan et al., 2016) to infer exact amplicon sequence variants (ASVs) instead of the rough and less precise 16S rRNA OTU clustering approach (Callahan, McMurdie, & Holmes, 2017; Dahan et al., 2018) that groups the sequences with a 97% identity (Edgar, 2018). First, we filtered by quality (trunQ = 25) and discarded the sequences if they presented more than two Ns (maxN = 2) or more than two expected errors (maxEE = 2). Next, the forward and reverse reads for each sample were combined into a single merged contig sequence, and we grouped all identical reads into unique sequences to determine their abundance. After building the ASVs table and removing chimeras (detected using self-referencing), sequences were classified and identified with *Decipher* v.2.26.0 (Wright, 2016), using the SILVA rRNA sequence database v.138 as the taxa reference (Quast et al., 2013). We used *phyloseq* v.1.42.0 (McMurdie & Holmes, 2013) to classify and remove any sequence not classified at the kingdom level or belonging to Archaea, Eukaria, chloroplasts, or mitochondria. For the last steps of the sequence cleaning process, we used *metagMisc* v.0.5.0 (Mikryukov, 2023) to eliminate ASVs with less than ten reads (minabund = 10). Additionally, we used *SourceTracker* (Knights et al., 2013) to estimate the proportion of the bacterial community in the blue whales blows samples that comes from the set of technical controls. ASVs detected in the LabControl sample were considered potential contaminants if their mean abundance was equivalent to or greater than 25% of their mean abundance in the whale samples (Caruso et al., 2019; Minich et al., 2019). Finally, we plotted Venn diagrams to show the logical relation of ASVs between blow and technical control samples using *MicEco* version 0.9.19 (Russel, 2023).

### Respiratory microbiome analysis

To get a sense of the community composition in the samples, we used *phyloseq* to identify the distribution of read counts from all the samples, and to plot rarefaction curves and the relative abundance stacked bar plot at the major level class and genus. Using *microbiome* v.1.20. (Lahti et al., 2017), we identified the common core bacteriome (threshold detection = 2/100, prevalence = 50/100). We selected these values because we wanted a more conservative approach and did not want to consider “rare bacteria” in the analysis. In addition, we analysed how the pattern of the common central microbiome changed based on a sliding prevalence range (10% - 100%) and a detection threshold of 0.02%. Using these criteria, we ran a linear regression model to determine the number of ASVs detected, given a detection threshold value for a specific prevalence. We calculated alpha diversity indices: richness (S) and Simpsońs diversity index (D) using *vegan* v.2.6.4 (Oksanen et al., 2019). All graphs were rendered using Tableau (Murray, 2013).

## Results

A total of 20 samples were collected and analysed in this study. These samples included 17 photo-identified blue whales and three technical controls. Exhaled breath samples were collected from animals in the Gulf of California using a small drone. No adverse behaviour was detected before, during, or after sampling (see Domínguez-Sánchez et al., 2018). After filtering, denoising, merging, and chimera elimination (2.33% of the reads), we recovered 72,254 sequences corresponding to 1,367 amplicon sequence variants (ASVs). Because of their low relative abundance, 677 of these ASVs were categorized as "others". Additionally, we eliminated 41 ASVs classified as Archea (n = 2), chloroplasts (n = 31), or mitochondria (n = 8), and 9 ASVs that were considered putative contaminants from LabControl. The sample coverage (i.e., the proportion of the total number of individuals in a community that belong to the species represented in the sample; (Chao & Chiu, 2016) exceeded 97.6% in all cases.

The logical relation analysis (Fig. 1) showed a total of 651 ASVs, of which 489 (76%) were identified solely in blow samples, 37 (6%) in SeaWater, 28 (4%) in HumanSneeze, and 11 (2%) in the LabControl sample. Alpha diversity measures revealed that species richness (S) in blow samples ranged from 70 to 440 (mean = 253.06) and Simpson’s index (D) ranged from 0.90 to 0.99 (mean = 0.97). The Compositional analysis identified 15 bacterial classes in the blow and technical control samples (Fig. 2), dominated by Gammaproteobacteria (27.14%), Bacteroidea (19.83%), and Clostridia (12.89%) as the most representative classes in the respiratory tract of blue whales. We also calculated the relative abundances of the 15 most representative bacterial genera in the blow samples (Fig. 3). *Psychrobacter* spp. (8.01%), *Oceanovirga* spp. (6.47%), *Tenacibaculum* spp. (5.92%), *Streptococcus* spp. (5.03%), and *Cutibacterium* spp. (4.49%) were the most abundant. Interestingly, two whale blows (Bm057 and Bm044) had a high relative abundance of opportunistic pathogens, *Mycoplasma* spp. (26.8%), and *Streptococcus* spp. (69.35%), respectively.

**Figure 1.**
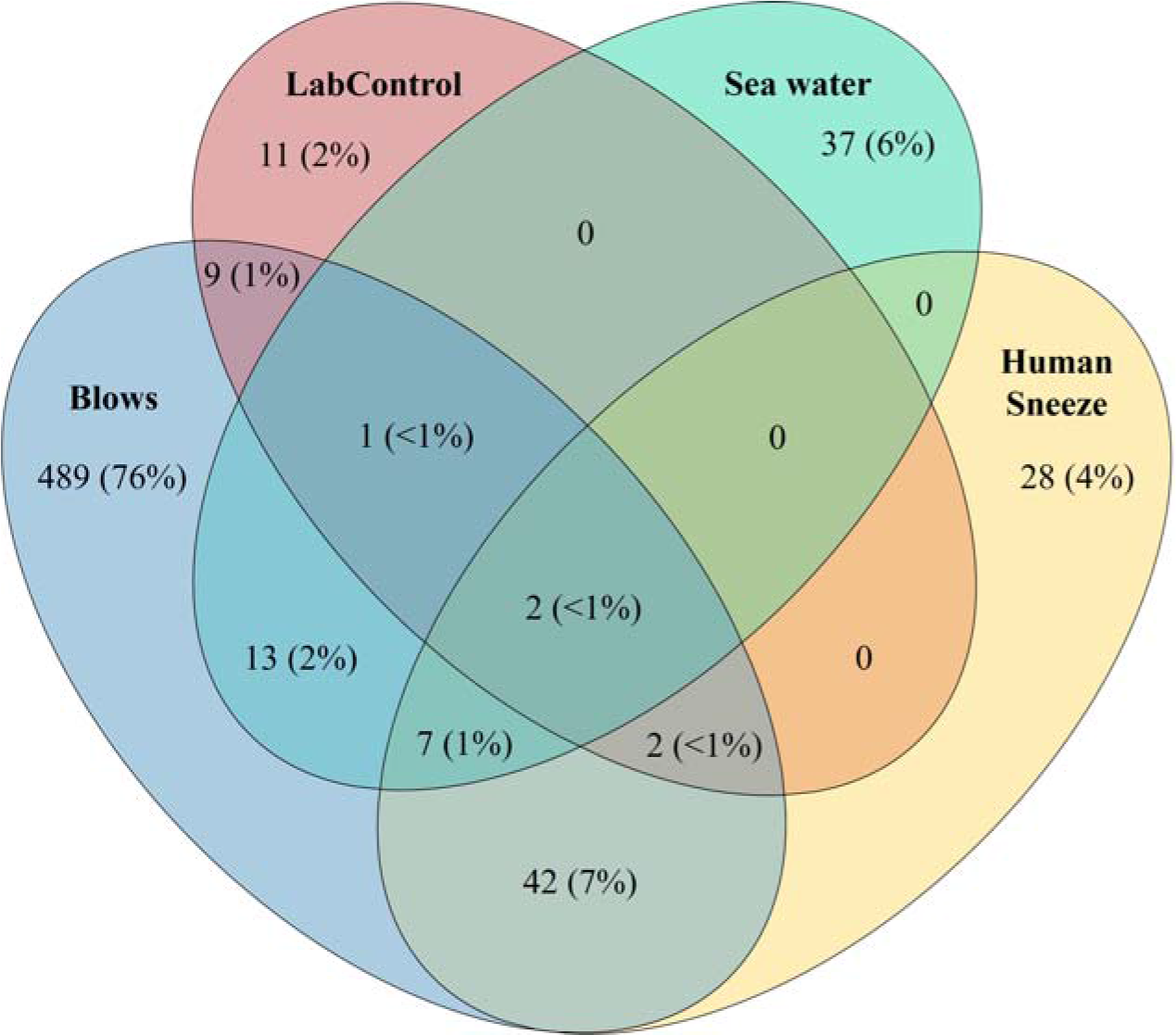
Representation of the ASVs shared among the Eastern North Pacific blue whale blow samples and technical control samples. The Venn diagram displays the ASV quantity shared among blows (blue), human sneeze (yellow), seawater (green) and Laboratory control (red).

**Figure 2.**
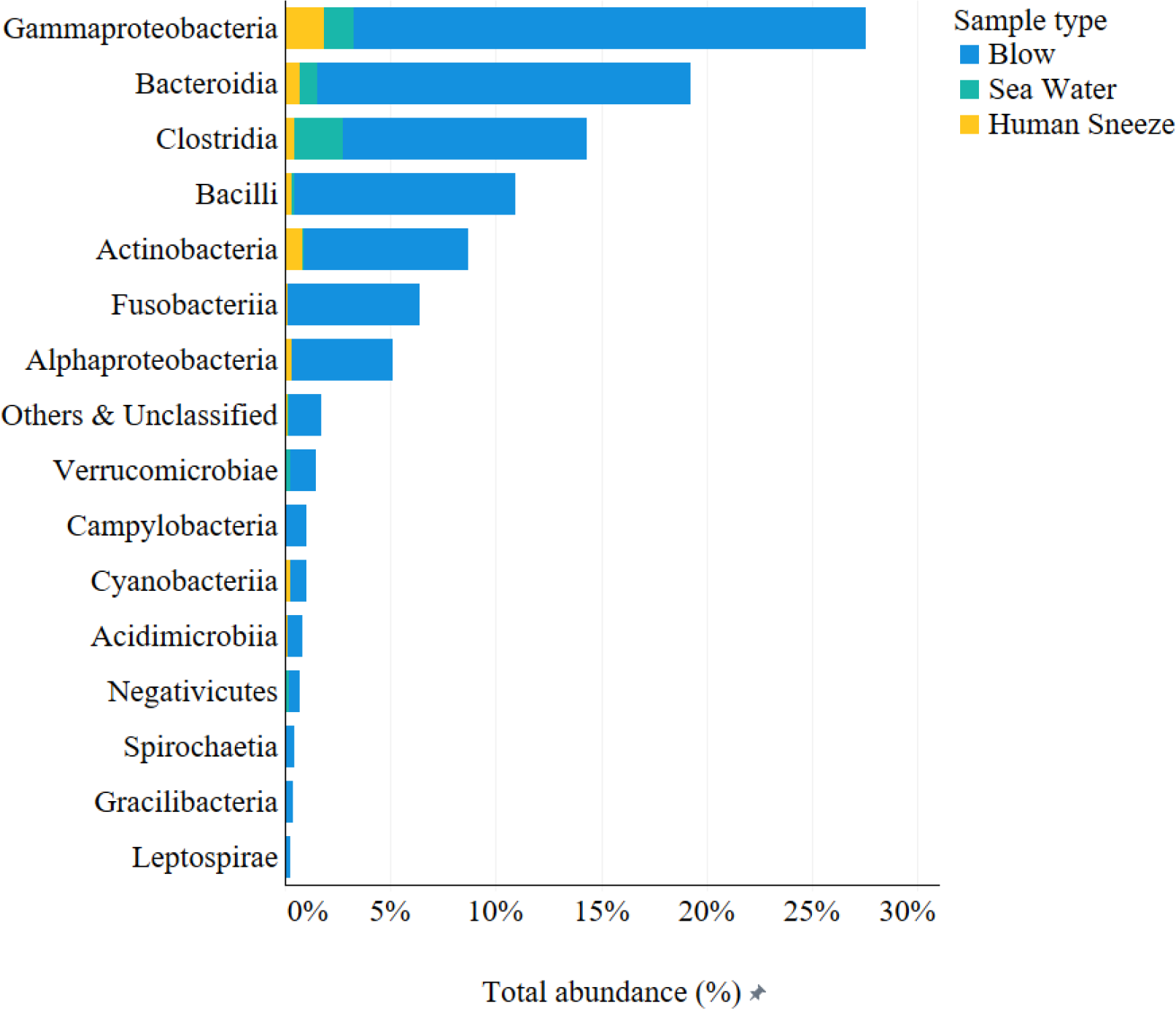
Total abundance (%) of the 15 bacterial classes identified in all samples. ASVs with less than 0.02% of relative abundance are grouped as “others” along with the unclassified.

**Figure 3.**
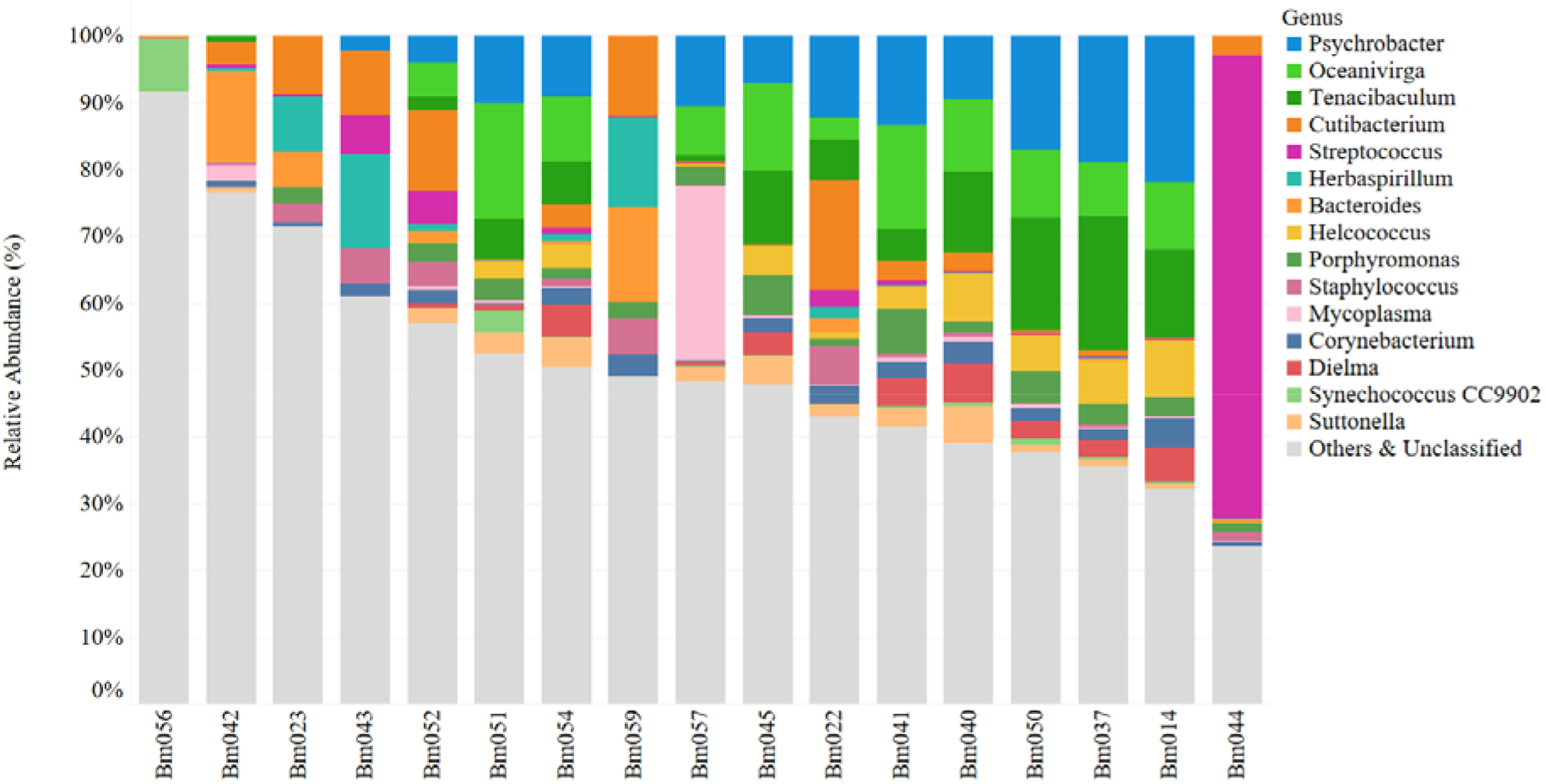
Stacked bar plot depicting relative abundance of the top 15 bacterial genera detected in the Eastern North Pacific blue whale blow samples. Each vertical bar depicts the relative abundance of adjusted sequence variants (ASVs) and associated taxa that were recovered per sample. Plot shows the top fifteen identified bacterial genera, unclassified are grouped along “others” (sum of bacteria that did not reach the detection threshold of 0.02%).

Eleven ASVs were considered the common core bacteriome of the respiratory tract of blue whales in the Gulf of California (Fig. 4a). The common core spanned nine classified ASVs in four bacterial genera (*Cutibacterium, Oceanivirga, Tenacibaculum*, and *Psychrobacter*) and two ASVs that were not classified at the genus level but belonged to the families Lachnospiraceae and Arcobacteraceae. Additionally, analysing changes in the pattern of the common central bacteriome, based on a range of prevalence and detection threshold values (Fig. 4b), it was possible to identify that the common central bacteriome of blue whale blow can vary from 232 ASVs (10% prevalence and 0.02 detection threshold) to 1 ASV (70% prevalence and 0.02 detection threshold), in all cases revealing *Cutibacterium* as the genus with the highest prevalence (70%) in blue whale blow samples. This bacterium was also identified in the technical controls, with a mean relative abundance of 4.73%. *Herbaspirillum* spp., the most abundant genus found in the seawater sample with 19.39% abundance, was also found in the blow samples with an average abundance of 2.38% and in three samples with higher abundances: Bm023 (8.21%), Bm043 (14.03%), and Bm059 (13.60%).

**Figure 4.**
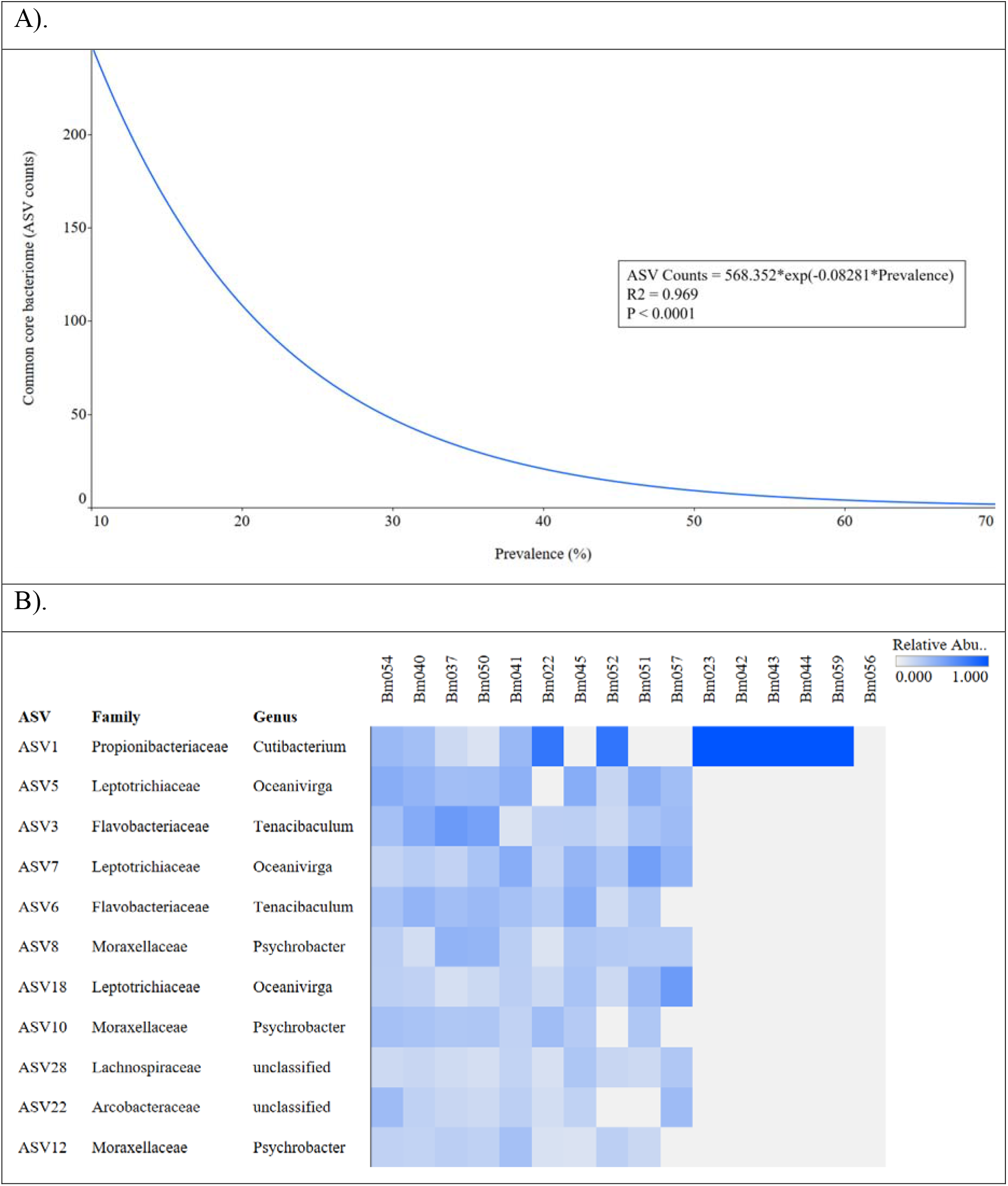
**a)** Regression model shows changes in the pattern of the common central respiratory bacteriome of the Eastern North Pacific blue whale, based on a range of prevalence and detection threshold values. **b)** Relative abundances of bacteria genera belonging to the core respiratory bacteriome of the blue whale (nine classified and two unclassified ASVs present in more than 50% of the samples and more than 0.02% of relative abundance).

## Discussion

Animal conservation can benefit from knowledge on disease and health status of threatened and endangered species (Heard et al., 2013; Pedersen et al., 2007). One of the factors that influence both disease and health of animals is the microbiome (Matthewman et al., 2023), and conservation science has recently begun to include studies of the microbiome to increase our understanding of population health (see Comizzoli & Power, 2019). Here, we examined the respiratory microbiome of the Eastern North Pacific blue whale, the planet’s largest extant animal and currently an endangered species for which there is extremely limited knowledge about their health and disease.

Recent studies of the human respiratory microbiome have shown that bacterial communities of the respiratory tract are key to maintaining respiratory health (Bond et al., 2017; Glendinning, McLachlan, & Vervelde, 2017; Watson, De Koff, & Bogaert, 2019), not only in terms of their metabolic contribution (Van Treuren & Dodd, 2020), but also because they prevent the colonization of the epithelium by environmental pathogens (Man, De Steenhuijsen Piters, & Bogaert, 2017; Yamamoto et al., 2021). Identifying the composition and abundance of the bacterial communities that constitute the microbiome of healthy individuals is an important step to establishing a baseline that will help identify bacteria associated with respiratory diseases (Lemon et al., 2010; N. Lima et al., 2012), assess chronic states of suboptimal health (Mackenzie et al., 2017) and predict community changes due to perturbation (Shade & Handelsman, 2012; Yamamoto et al., 2021)

The results of our study demonstrate that the blow of the Eastern North Pacific blue whale supports a diverse and rich bacterial community. We identified 1,367 ASVs with high sample coverage and varying levels of richness and relative abundance among samples. This could be an indicator of temporary fluctuations in the composition of the microbiome (Eloe-fadrosh & Rasko, 2013). The richness and relative abundance of the microbiome vary among healthy animals (Coyte et al., 2019; Toro et al., 2021), due mainly to bacterial immigration from the environment to the lungs during inhalation, bacterial elimination via mucociliary clearance, and a relatively small contribution of the growth rate of each bacterial community (Dickson, Erb-Downward, & Huffnagle, 2014; Dickson et al., 2016; Huffnagle, Dickson, & Lukacs, 2017). Evidently, we cannot rule out that the variation in richness and relative abundance that we observed was also related to differences in the volume of blow collected, which due to the nature of the samples collection technique used, could not be standardized. On one hand, whales are likely to differ in the amount of blow exhaled, depending on the whale’s size, and the depth and duration of the dive. Furthermore, although drones are safe, minimally invasive, and do not seem to affect the whales during blow collection (Domínguez-Sánchez et al., 2018), inherent limitations such as flight height and different wind conditions, could potentially result in different volumes of blow being collected (Apprill et al., 2017).

Various indices have been used to estimate the diversity of microbial communities. In our study, we used Simpson’s diversity index because it considers both richness and evenness (Johnson & Burnet, 2016) and was previously identified as one of the most accurate estimators of diversity in an unknown bacterial community (Haegeman et al., 2014). In our study, Simpson’s diversity index averaged 0.97 (minimum = 0.90, maximum = 0.99), demonstrating high bacterial diversity in the blow of the whales that we sampled. Some studies have shown that the microbiome of a healthy animal tends to have high diversity, which presumably allows it to tolerate or counteract changes arising from extrinsic challenges (Chan, Estaki, & Gibson, 2013; Van Cise et al., 2020). Bacterial diversity was high in nearly all of the blue whale blow samples, and the composition of the microbiome was dominated mainly by members of the phyla Proteobacteria, Firmicutes, Bacteroidota, Actinobacteria, and Fusobacteriota, which have been reported to be major components of the healthy respiratory microbiome of other mammals (Chaban et al., 2013; Rhodes et al., 2022). Simpsońs diversity index was similar to that reported for humpback whale blows (Apprill et al., 2017) and bottlenose dolphin blowholes (Johnson et al., 2009b; Bik et al., 2016). It is likely that these results are evidence that the respiratory microbiome diversity (based on Simpsońs index) is conserved among cetaceans. However, the taxonomic richness of the blue whale blow was nearly twice that reported for humpback whales. In our samples, the percentage of "others" (sum of bacteria that did not reach the detection threshold of 0.02%) was higher than that in humpback whales (Apprill et al., 2017). Based on this result, we suggest that the microbiome of the Eastern North Pacific blue whale respiratory tract may be more complex than that of humpback whales. At this stage we can only speculate about the reasons that could explain such a difference. They may be due to the use of DADA2 for resolving ASVs rather than minimum entropy decomposition (MEDs; Eren et al., 2015), which has been used previously, given that although similar in what both approaches resolve, ASVs are better at removing erroneous sequences (Callahan et al., 2016; Ahlgren et al., 2019). As the algorithm DADA2 allows for the independent analysis or grouping of samples, we conducted pooled analyses to increase the sensitivity to detect ASVs that could be present at very low frequencies in multiple samples (Callahan et al., 2016). This approach could explain why a higher percentage (mean = 30.91%) of “rare bacterial biosphere” (Pedrós-Alió, 2012) was identified in our study than in the humpback whale study (see (Apprill et al., 2017). This “rare bacterial biosphere”, formed by bacteria that are present at low relative abundances, is particularly important for dealing with dysbiosis, as they could be considered as a seed bank of genetic resources that can lead to the restoration of the core microbiome (Pedrós-Alió, 2012; Skopina et al., 2016; Jousset et al., 2017). A more complex respiratory microbial community is likely to be beneficial to whales, given that microbiomes with a higher richness of species have more synergistic interactions between bacterial taxa, which improve the functioning of the ecosystem (Bell et al., 2005). It is worth mentioning that most studies published to date do not consider the bacterial taxa found in low abundances to be relevant; however, these small populations are now thought to play an important role for the functioning of the ecosystem (Willis, Bunge, & Whitman, 2017) and host health (Jousset et al., 2017). It has been demonstrated that a high diversity of low-abundance bacteria is correlated with less severe bacterial infections in human lungs (Van Der Gast et al., 2010).

Sixteen bacterial classes were identified in the blow samples, some of which were also found in the seawater samples. This is unsurprising, as there is likely to be some seawater carried over when whales exhale. However, the blow samples harboured ASVs belonging to bacterial genera that were not found in the seawater sample, such as *Psychrobacter, Oceanivirga, Tenacibaculum, Helcococcus, Porphyromonas, Mycoplasma, Dielma, Synechococcus,* and *Suttonella*. This shows that despite the potential carryover of seawater to the blow during exhalation, the bacterial communities of the blue whale respiratory tract are different from those of seawater.

Eleven ASVs belonging to the families Propionibacteriacea, Flavobacteriaceae, Lachnospiraceae, Leptotrichiaceae, Moraxellaceae, Porphyromonadaceae, and Arcobacteraceae; were shared in more than half of the blue whale samples and were considered to be the common core respiratory bacteriome of the Eastern North Pacific blue whale. The interindividual variability of the respiratory microbiome appears to be higher than that of the humpback whales, as 25 distinct bacteria were found to be shared among all the animals sampled (Bierlich et al., 2018). *Psychrobacter* spp. and *Tenacibaculum* spp. have been reported to be key members of the core microbiome in other baleen whales (Hooper, Littman, & Macpherson, 2012; Bierlich et al., 2018; Van Cise et al., 2020; Toro et al., 2021). *Porphyromonas* spp. have previously been isolated from humpback whale skin (Apprill et al., 2014) and humpback whale blows (Apprill et al., 2017). Furthermore, *Porphyromonas* spp. and *Fusobacterium* spp. have been described as bacteria of the core pulmonary microbiome in humans (Erb-Downward et al., 2011; Charlson et al., 2012; Cui et al., 2014; Huang, Yang, & Li, 2016). These bacteria have also been reported sporadically and in low abundance in the sheep respiratory tract (Glendinning et al., 2016). We also identified *Moraxella* spp. in blue whale blows. This bacterial genus is present in humpback whale blows (Apprill et al., 2017) and is commonly found in the lungs of healthy dogs (Tress et al., 2017). However, it has also been reported in humans (Yi et al., 2014) and cattle with respiratory diseases (Lima et al., 2016). The bacteria identified in the blue whale respiratory tract are similar to those reported in other cetaceans and terrestrial mammals, and some are known to cause diseases. At this stage, we were unable to unequivocally establish that the health of the whales sampled was not compromised; however, given that they were present in most of the whales, we can assume that they are part of their respiratory microbiome and that they are likely to reflect a healthy respiratory tract.

Interestingly, five bacterial genera (*Psychrobacter, Tenacibaculum, Staphylococcus, Cutinebacterium,* and *Corinebacterium*) identified in the blue whale blow are associated with the skin of humans and other terrestrial mammals (Grice & Segre, 2011; Byrd, Belkaid, & Segre, 2018; Worthing et al., 2018) and were recently identified as part of the skin microbiota of captive bottlenose dolphins *Tursiops truncatus*, killer whales *Orcinus orca*, and free-ranging humpback whales (Hooper, Littman, & Macpherson, 2012; Apprill et al., 2014; Chiarello et al., 2017; Rhodes et al., 2022). It is likely that their presence in the blow samples indicates that they colonize the blowhole epithelial lining of blue whales and are forcefully expelled during exhalation (Apprill et al., 2017), leading to their presence in the blow samples.

*Mycoplasma* spp. was detected in one sample (Bm057) with a 26.84% relative abundance. The presence of these bacteria could be indicative of a transient bloom within the respiratory tract or an active respiratory infection, considering that these bacteria are typically present in the respiratory tract of mammals at a low abundance, but during active pathological processes, such as pneumonia and other respiratory conditions, their relative abundance increases (Dai et al., 2018). In marine mammals, *Mycoplasma* spp. has been associated with respiratory disease and has been detected in the lungs of stranded harbour porpoises *Phocoena phocoena*, Sowerby’s beaked whales *Mesoplodon bidens* (Foster et al., 2011) and California sea lions *Zalophus californianus* (Haulena et al., 2013) during unusual mortality events. Furthermore, although this genus has been found in healthy killer whales (Raverty et al., 2017; Rhodes et al., 2022), the role of *Mycoplasma* spp. during disease episodes in cetaceans, as well as their host specificity and association with cetacean stranding events, remains poorly understood (Foster et al., 2011; Rhodes et al., 2022). It is certainly possible that the whale from which sample Bm057 was collected was experiencing a respiratory infection that involved *Mycoplasma* spp. Gaining clinical information that would allow us to establish this beyond any doubt is not feasible, but we propose that future studies of blue whale populations consider the presence of *Mycoplasma* spp. as an indicator of suboptimal respiratory health.

Infectious diseases are one of the leading causes of death in marine mammals (Numberger et al., 2021; Morick et al., 2022), and *Streptococcus* spp. is one of the most reported pathogens of pinnipeds and cetaceans (Bianchi et al., 2018). We identified one sample (Bm044) with a high relative abundance (69.35%) of *Streptococcus* spp. Overall, ten streptococcal species have been isolated and identified multiple times from twenty-three pinniped and cetacean species worldwide (Numberger et al., 2021). *Streptococcus agalactiae*, for example, was isolated from a wound and navel infection in grey seals *Halichoerus grapus* (Baker, 1988), as well as lung and spleen samples from Antarctic fur seals *Arctocephalus gazella* (Baker & McCann, 1989). *Streptococcus phocae* is frequently found in lung lesions, implying commensal colonization of the oropharynx and subsequent opportunistic respiratory tract infections (Taurisano et al., 2018), and *Streptococcus iniae* is a significant aquatic pathogen of farmed fish species, an important zoonotic pathogen, and an identified cause of disease in captive Amazon River dolphins *Inia geoffrensis* (Souter et al., 2021), bottlenose dolphins (Evans et al., 2006), and death in short-beaked common dolphins (Souter et al., 2021).

Having detected two potentially pathogenic bacterial genera in this study could mean that the Eastern North Pacific blue whale is not commonly in contact with coastal areas where spillover of pathogens from humans or domestic animals could occur. This would be in stark contrast to killer whales, which live in areas where there are a large number of environmental anthropic stressors (Raverty et al., 2017). However, a previous study reported *Entamoeba* spp., *Giardia* spp., and *Balantidium* spp., most likely from sewage discharge, in faeces of blue whales from the Gulf of California (Pacheco-Armenta, 2019). Therefore, it is plausible that rather than limited exposure, the presence of *Mycoplasma* spp. and *Streptococcus* spp. in two individuals reflect a suboptimal immune status or an underlying upper or lower respiratory condition, which could allow respiratory colonization of this pathogen.

It is important to note that the respiratory microbiome of the blue whales analysed in our study harboured bacteria that are commonly found in the oropharynx, nasopharynx, and mouth of different terrestrial mammals (German & Palmer, 2006; Guglielmetti et al., 2010), in which those anatomical structures are interconnected in the upper respiratory tract. In contrast, cetaceans have no anatomical connection between the nasopharynx and mouth (Apprill et al., 2017; Smith, Tang, & Reidenberg, 2017). This finding provides strong evidence that the core microbiome that we described belongs to the respiratory system of blue whales and does not include their oral bacteria.

Given the current state of our oceans, which face habitat degradation, pollution, and other anthropogenic stressors (Melcón et al., 2012; Mouton & Botha, 2012; Palmer et al., 2022); suboptimal immune responses can occur in top-predator marine animals (Acevedo-Whitehouse & Duffus, 2009; Van Bressem et al., 2014; Hall et al., 2018), in turn increasing the risk of diseases in their populations (Sós et al., 2013; Van Bressem et al., 2015; Reisfeld et al., 2019) The study of the respiratory bacteriome of blue whales provides an excellent opportunity to develop conservation plans for this endangered species. Researchers can develop effective strategies to protect blue whales from disease, pollution, and other threats by better understanding the microbial communities inhabiting their respiratory tracts. To use the respiratory bacteriome as a tool to help assess the health of large whales (Apprill et al., 2017; Toro et al., 2021; Rhodes et al., 2022), it is imperative to first identify the composition of the microbiome of the respiratory tract of different species, which is what we have done for the Eastern North Pacific blue whales.

## Conclusion

To the best of our knowledge, this is the first study to characterize the microbiome of the respiratory tract of blue whales. We found that the Eastern North Pacific blue whales sampled in the Gulf of California harboured a similar respiratory bacterial composition among individuals. Additionally, richness and relative abundance results were comparable with those reported in the microbiome of healthy animals and humans; therefore, we propose that the core respiratory bacteriome identified here could be used as a baseline for future long-term studies aimed at identifying shifts in the composition and co-occurrence patterns of the respiratory microbiome and identify ASVs related to changes in body condition, as a proxy for poor health condition. This knowledge can help conservation efforts to protect blue whales during their long migrations and ensure that future generations will continue to marvel these incredible creatures and appreciate their ecological role in maintaining the health of our oceans.

### Ethical approval and consent to participate

This study complied with the recommendations and methods for approaching blue whales provided by Mexican legislation (NOM-059-SEMARNAT-2010). All procedures were approved by the Bioethics Committee of the Universidad Autónoma de Queretaro (Mexico), and sampling was conducted under permits SGPA/DGVS/00255/16 and SGPA/DGVS/01832/17 issued by the Dirección General de Vida Silvestre to D. Gendron.

## Acknowledgements

We thank Manuel Antonio Zamarrón Nunez for his assistance during navigation, and Ana Sofia Merino, Aurora Paniagua, Madeleine Gauthier, Daniel Valdivia and Ricardo Mirsha Mata Cruz for their help during fieldwork. CADS was funded by a CONACYT PhD Studentship (558253). Fieldwork (sampling and navigation) was funded by The Instituto Politécnico Nacional (SIP20160496 and 2017014), Rufford Foundation 2nd small grant for Nature Conservation (2017), and the Program for the Conservation of Species at Risk (Programa de Conservación de Especies en Riesgo, Comisión Nacional de Áreas Naturales Protegidas). Molecular analysis was partly financed by a Small Grant in Aid of Research from the Society for Marine Mammalogy.

## Availability of data and materials

The datasets used and analysed during the current study are available from the corresponding author upon reasonable request.

## Authors’ contributions

CAD collected the samples, performed molecular analyses, analysed the data, and drafted the manuscript. RCA conducted statistical programming for microbiome analysis and network construction and helped interpret the results. DG conducted fieldwork, collected samples, and co-supervised the research. KAW conceived, designed, and supervised the research. All authors read and commented the final draft of the manuscript and gave approval for publication.

